# First evidence of virus-like particles in the bacterial symbionts of Bryozoa

**DOI:** 10.1101/2020.04.16.045880

**Authors:** A.E. Vishnyakov, N.P. Karagodina, G. Lim-Fong, P.A. Ivanov, T.F. Schwaha, A.V. Letarov, A.N. Ostrovsky

## Abstract

Bacteriophage communities associated with humans and vertebrate animals have been extensively studied, but the data on phages living in invertebrates remain scarce. In fact, they have never been reported for most animal phyla. Our ultrastructural study showed for the first time a variety of virus-like particles (VLPs) and supposed virus-related structures inside symbiotic bacteria in two marine species from the phylum Bryozoa, the cheilostomes *Bugula neritina* and *Paralicornia sinuosa*. We also documented the effect of VLPs on bacterial hosts: we explain different bacterial ‘ultrastructural types’ detected in bryozoan tissues as stages in the gradual destruction of prokaryotic cells caused by viral multiplication during the lytic cycle. We speculate that viruses destroying bacteria regulate symbiont numbers in the bryozoan hosts, a phenomenon known in some insects. We develop two hypotheses explaining exo- and endogenous circulation of the viruses during the life-cycle of *B. neritina*. Finally, we compare unusual ‘sea-urchin’-like structures found in the collapsed bacteria in *P. sinuosa* with so-called metamorphosis associated complexes (MACs) known to trigger larval metamorphosis in a polychaete worm.

**Importance:** Complex symbiotic systems, including metazoan hosts, their bacterial symbionts and bacteriophages are widely studied using vertebrate models whereas much less is known about invertebrates. Our ultrastructural research revealed replication of the viruses and/or activation of virus related elements in the bacterial symbionts inhabiting tissues of the marine colonial invertebrates (phylum Bryozoa). The virus activity in the bacterial cells that are believed to be transmitted exclusively vertically is of a special importance. In addition, in the bacterial symbionts of one of the bryozoan hosts we observed the massive replication of the structures seemingly related to the Metamorphosis associated complexes (MAC). To our knowledge, MACs were never reported in the animal prokaryotic symbionts. Our findings indicate that Bryozoa may be new suitable model to study the role of bacteriophages and phage-related structures in the complex symbiotic systems hosted by marine invertebrates.

## INTRODUCTION

Viruses are found in all kingdoms of living organisms and are best studied in those that have an applied or medical value (Ohmann & Babiuk 1986; Woolhouse et al. 2012; Johnson et al. 2015; Glennon et al. 2018; https://talk.ictvonline.org/taxonomy/). In addition to harboring the viruses replicating in eukaryotic cells, all known animals (as well as other multicellular organisms) are associated with specific microbial communities that include viruses infecting their symbiotic microorganisms. Most of these viruses are bacteriophages. Although the bacteriophage communities (viromes) of vertebrates are much better studied (reviewed in Letarov and Kulikov 2009; Shkoporov 2019; Kwok et al. 2020), some data are also available for invertebrates. These include the viral communities associated with the cnidarian *Hydra* (Grasis et al. 2014; Bosch et al. 2015) and certain scleractinian corals (Weynberg et al. 2017; Mahmoud and Jose 2017), which have been recently characterized using metagenomic methods. These studies showed that such communities are species-specific and may be significant for homeostasis of the animal hosts.

In addition to the complex microbiomes associated with invertebrate digestive systems or body surfaces, some species harbor specific bacterial symbionts that may be vertically transmitted. Although such symbionts are either intracellular or live in the host tissues (sometimes in special organs), they can also harbor bacteriophages. The best-known example is *Wollbachia* bacteriophage WO. The symbiotic bacteria *Wolbachia* exert a major influence on their arthropod hosts by manipulating their reproduction and providing increased resistance to infections (López-Madrigal and Duarte 2019). In turn, the bacteriophage WO influences the bacterial titer in the host tissues (Bordenstein et al. 2006). This phage also encodes proteins important for bacterium interactions with the host (Perlmutter et al. 2019).

In the marine realm, the large variety of viruses are found across diverse taxa, including protists and various invertebrates such as sponges, cnidarians, flatworms, polychaetes, mollusks, crustaceans and echinoderms (reviewed in Johnson 1984; Weinbauer 2004; Munn 2006; Lang et al. 2009; Rosario et al. 2015; see also Reuter 1975; Vijayan et al. 2005; Nobiron et al. 2008; Crespo-González et al. 2008; Marhaver et al. 2008; Claverie et al. 2009; Jackson et al. 2016 and references therein). Being present in large numbers in the seawater (Suttle 2005; Brum et al. 2013), viruses, among others, enter suspension-feeders (e.g. Middelboe & Brussaard 2017). Indeed, by filtering enormous volumes of water, sea sponges acquire viruses that infect their cells (Vacelet & Gallissian 1978; Luter et al. 2010; Pascelli et al. 2018) as well as their bacterial symbionts (Lohr et al. 2005). Corals, together with their eukaryotic and prokaryotic symbionts, harbor a variety of viruses too (e.g. Lohr et al. 2007; Patten et al. 2008; van Oppen et al. 2009; Vega Thurber & Correa 2011; Leruste et al. 2012; Pollock et al. 2014; Correa et al. 2016). Similarly, filter-feeders such as bivalve mollusks and tunicates and bacteria living in them also acquire viruses (reviewed in Farley 1978; Elston 1997; Renault & Novoa 2004; Richards et al., 2019). Recently a bacteriophage was found to enhance the biofilm formation in the gut of an ascidian by interacting with its bacterial hosts (Leigh et al. 2017).

The phylum Bryozoa is comprised of active filterers that feed mainly on microscopic algae, gathering them out of seawater (Winston 1977, 1978; Shunatova & Ostrovsky 2001, 2002; Schwaha et al. 2020). Together with sponges and cnidarians this group of colonial invertebrates is among the dominating foulers in many bottom communities from the intertidal zone to a depth of 8 km (Ryland 1970, 2005; McKinney & Jackson 1989; Nielsen 2013). Although viruses were never reported from bryozoans before, symbiotic associations with bacteria are known in species from several families of the order Cheilostomata, the largest bryozoan group (e.g. Lutaud 1965, 1969, 1986; Woollacott & Zimmer 1975; Dyrynda & King 1982; Moosbrugger et al. 2012; Mathew et al. 2018, reviewed in Karagodina et al. 2018). The symbionts are vertically transmitted via the larval stage (Woollacott 1981; Zimmer & Woollacott 1983, 1989; Boyle et al. 1987; Lim & Haygood 2004; Sharp et al. 2007a; Lim-Fong et al. 2008, and references therein). During our ongoing research we discovered presumed virus-like particles and virus-related structures associated with bacteria in two bryozoan species from two different families and distant localities. This paper presents the first description of the VLP (supposedly bacteriophages) in Bryozoa and their presumed effect onto their bacterial hosts. We also discuss the possible ways of the virus transmission and circulation and their role in these symbiotic systems.

## MATERIALS AND METHODS

### Animal material collection, fixation and microscopy

Colonies of the cheilostome bryozoan *Bugula neritina* (Linnaeus, 1758) (Bugulidae) (Fig. 1A) were collected intertidally on Atlantic Beach, Jaycee Park, Morehead City, North Carolina, USA, in spring 2011. *Paralicornia sinuosa* (Canu and Bassler, 1927) (Candidae) (Fig. 1B) was collected by SCUBA-diving in the vicinity of the Lizard Island Research Station, Great Barrier Reef, Coral Sea, Australia, on 4, 5 and 10 October 2012 between 6 and 12 m depth.

**Fig. 1.**
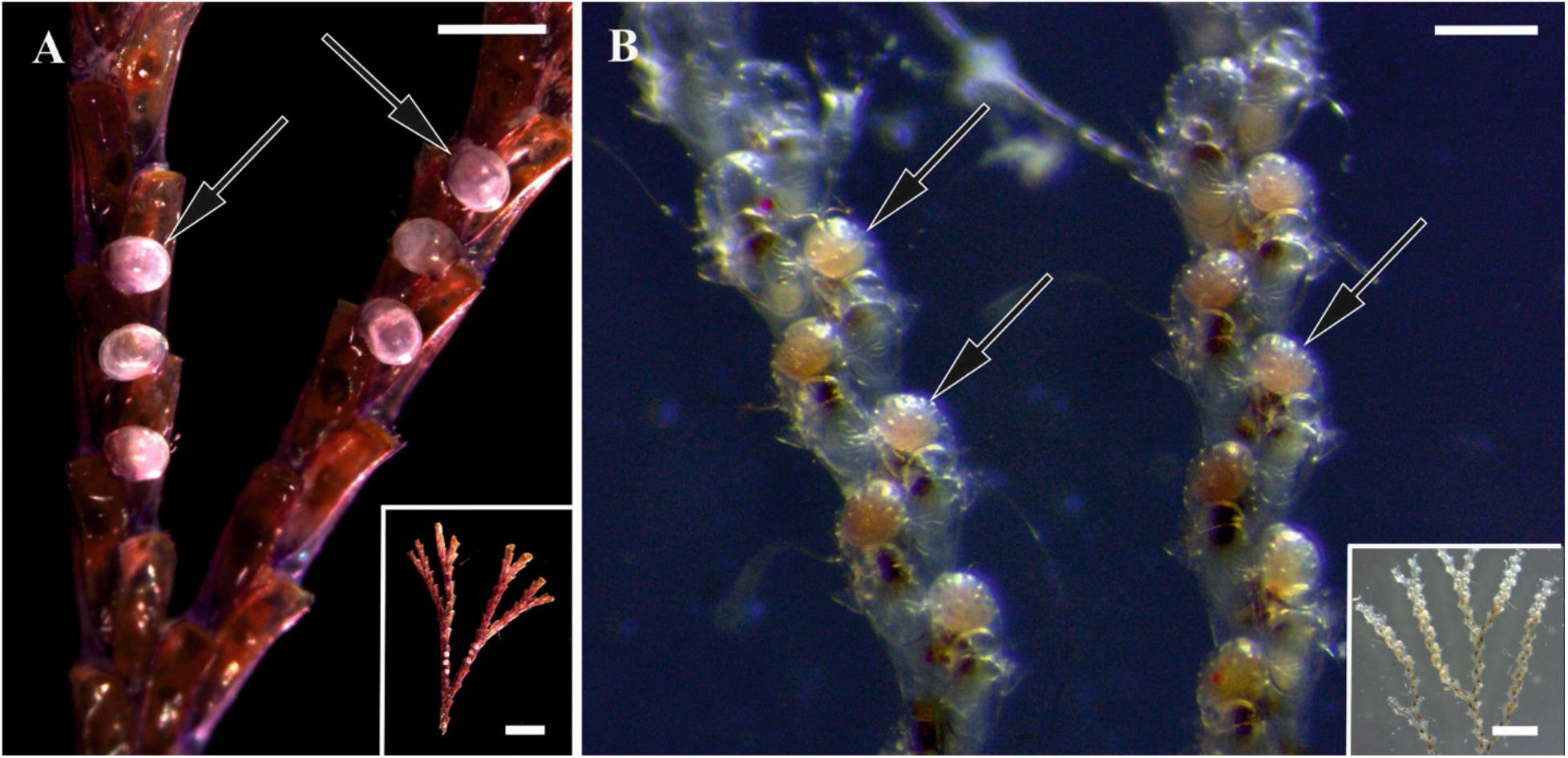
General view of colony branches of (A, insert) *Bugula neritina* and (B, insert) *Paralicornia sinuosa*. Arrows: brood chambers (ovicells) with embryos. Stereomicroscope. Scale bars: A, 500 μm, insert, 2 mm; B, 200 μm, insert, 1 mm.

Colony fragments were fixed in 2.5% glutaraldehyde (buffered in 0.1M Na-cacodylate containing 10% sucrose (pH 7.4)). They were postfixed with a 1 % solution of osmium tetroxide (OsO_4_). Decalcification was conducted for 24 h in 2% aqueous solution of EDTA. After this step the fragments were dehydrated in a graded ethanol series (30-50-70-80-90-100%), embedded in epoxy resin type TAAB 812 and sectioned (70 nm thick) using a Leica EM UC7 microtome (Leica Microsystems, Wetzlar, Germany). To find the area for transmission electron microscopy (TEM), the histological sections (1.0 μm thick) were prepared for light microscopy and stained with Richardson’s stain using standard methods (Richardson et al. 1960). We sectioned seven branches of *B. neritina* and four branches of *P. sinuosa*. Ultrathin sections were picked up with single slot copper grids with formvar support film and contrasted with uranyl acetate and lead citrate. Semithin sections were analyzed using an AxioImager.A1, Zeiss microscope (Zeiss, Oberkochen, Germany). Ultrathin sections were examined with a Jeol JEM-1400 microscope (JEOL Ltd., Japan).

### Bacterial and bacteriophage strains and their cultivation

To compare the objects found in the bacterial symbionts of *Bugula neritina* and reminiscent the VLPs, we prepared a suspension of *Escherichia coli* cells infected by bacteriophage RB49 (a T4-related virus) and visualized them using the same fixative and following the same procedure as used for bryozoan samples.

*E. coli* C600 strain and bacteriophage RB49 were from the collection of the Laboratory of Microbial Viruses, Winogradsky Institute of Microbiology. For the infection experiment the overnight culture of *E. coli* F5 (10.1007/s00705-019-04371-1) was grown in LB medium at 37°C with agitation. The culture was diluted 100-fold with the same medium and cultured under the same conditions up to OD_600_ 0.3. This optical density corresponds to 2×10^7^ c.f.u. ml^−1^. Phage RB49 lysate was added to 3 ml of the bacterial culture up to the multiplicity of infection of 5. The culture was further incubated 15 min and then the cells from 1 ml of the infected culture were spun down in the table-top centrifuge at 10 000 g for 40 seconds. The supernatant was removed and 200 μl of the same fixative solution that was used for the animal material was added, and the samples were then processed for thin sectioning and TEM study following the same protocol.

## RESULTS

Colonies of both studied species, *Bugula neritina* and *Paralicornia sinuosa*, are lightly calcified, erect and branched (Fig. 1). Histological sections and TEM study of colony fragments showed the presence of symbiotic bacteria in the funicular system in both species (Fig. 2). In *B. neritina* the densely packed bacterial cells were present in so-called ‘funicular bodies’, which are swollen areas of the transport funicular cords crossing the coelomic cavity of autozooids (Fig. 2A, B), and in *P. sinuosa* they were present inside the funicular cords themselves (Fig. 2C, D). Also, in *B. neritina*, bacteria were additionally recorded in and between the epidermal cells of the tentacles, in the cells of the body wall (epithelium of the introvert and peritoneal cells) and in the presumed amoebocytes situated on the epithelial lining of the ooecial vesicle plugging an entrance to the brood chamber (ovicell). In both species we detected intact (non-modified) as well as morphologically altered bacterial cells.

**Fig. 2.**
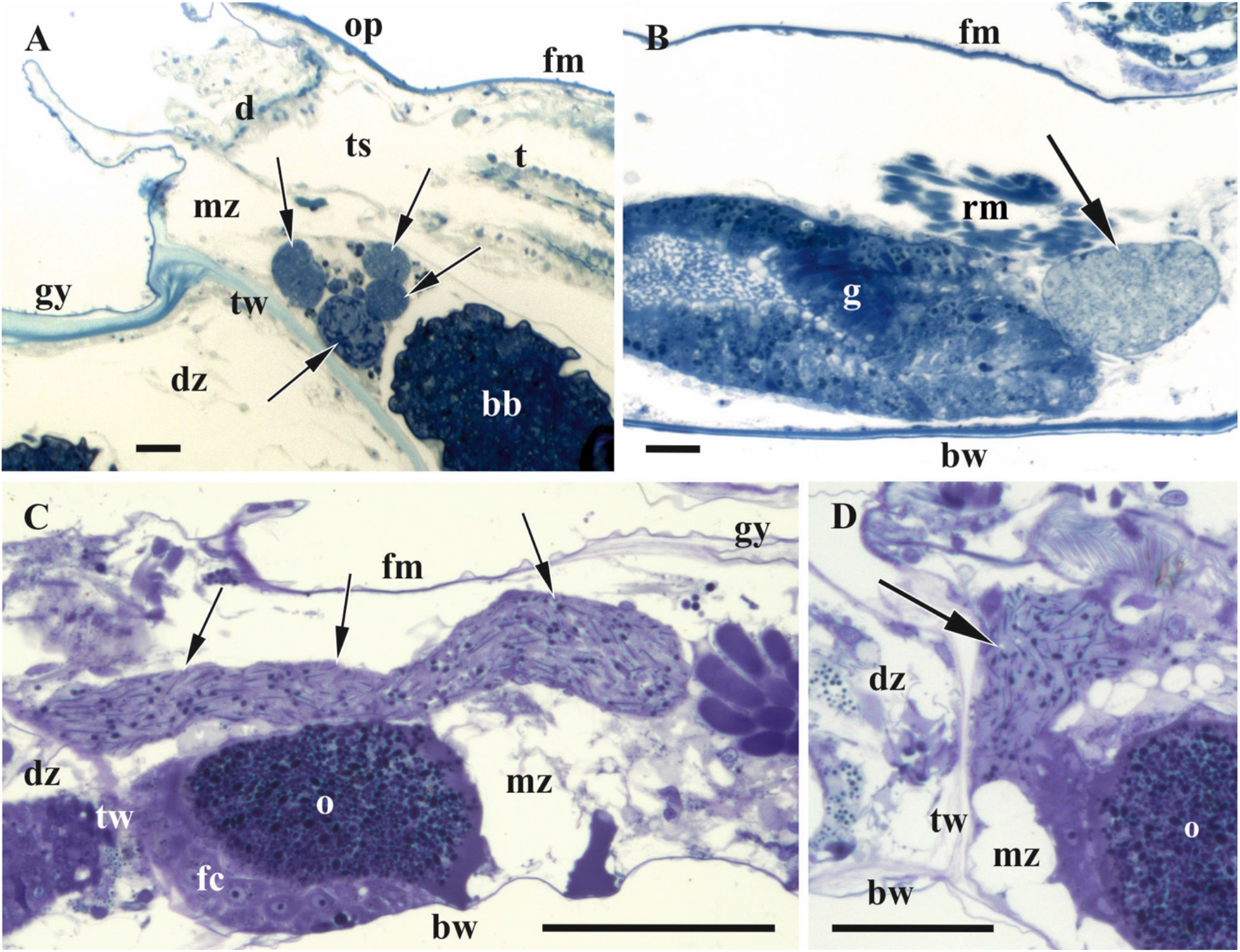
Histological sections of autozooids of (A, B) *Bugula neritina* and (C, D) *Paralicornia sinuosa* containing bacteria inside funicular bodies (A, B) or funicular cords (C, D) (shown by arrows). Paracrystalline structures are visible as black ‘dots’ and ‘lines’ inside funicular cords in C and D. Light microscopy. Abbreviations: bb, brown body (degenerated polypide); bw, basal wall; d, diaphragm; dz, distal zooid; fc, follicle cells; fm, frontal membrane; g, gut; gy, gymnocyst; mz, maternal zooid; o, oocyte inside ovary; op, operculum; rm, retractor muscles; t, tentacles of retracted polypide; ts, tentacle sheath; tw, transverse wall. Scale bars: A, B, 20 μm; C, 500 μm; D, 300 μm.

Ultrastructural study of the bacterial symbionts in both bryozoan hosts showed the presence of objects resembling virus-like particles (VLP) and/or virus-related structures. Our interpretation of the discovered particles (see below) as VLPs was based on their size, morphological features and in their occurrence in/near structurally altered bacterial cells. Although the virions of bacteriophages are dimensionally and morphologically stable and uniform, both fixation and TEM observations potentially could generate a ‘false’ visual diversity since the capsids are sectioned at different levels and viewed at different angles, and the tails (if present) may be hidden.

Comparison of *E. coli* cells infected by the phage RB49 showed that the bacteriophage heads and proheads exhibited a significant degree of apparent morphological variation (Fig. 3) similar to VLPs observed in bryozoans (Figs. 4, 6–8). The tails of RB49 particles could not be reliably seen inside the infected bacterial cells.

**Fig. 3.**
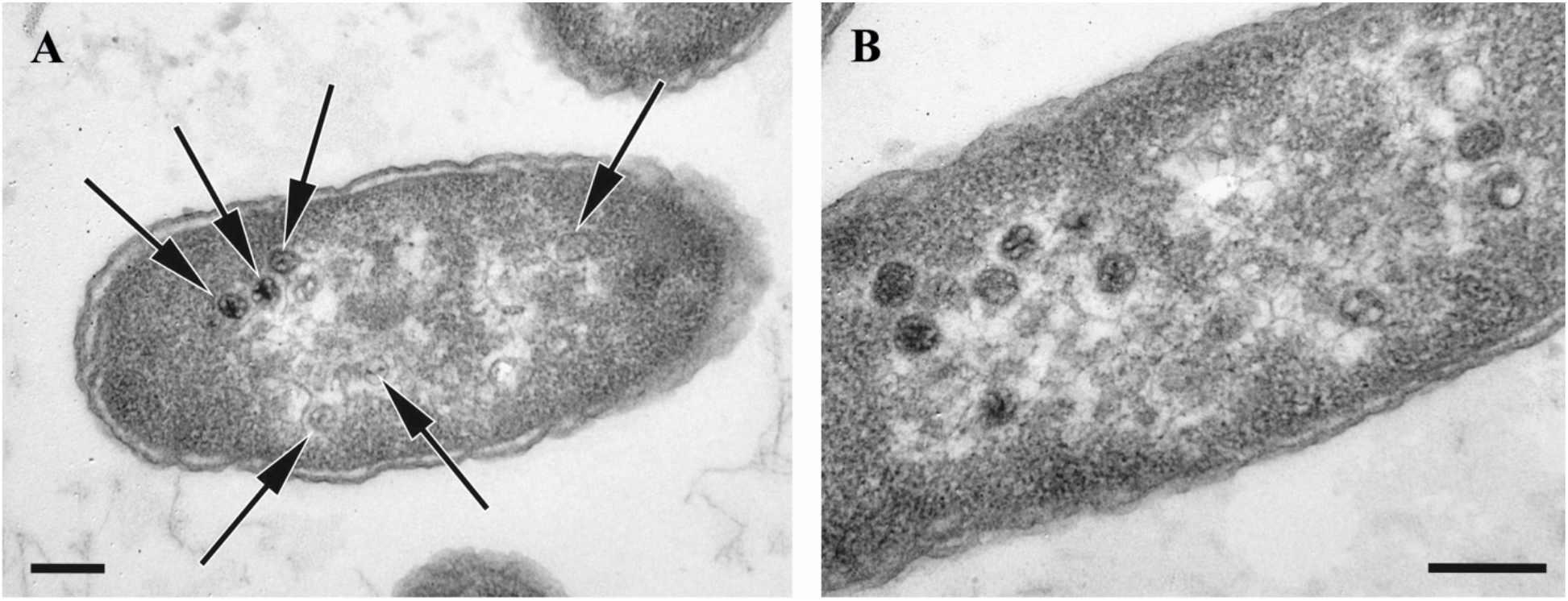
Thin sections of *E. coli* F5 cells infected by the bacteriophage RB49 (15 min post infection). Bacteriophage heads and proheads (both empty and partially filled) are visible (arrows on A). Note that the capsids do not appear exactly uniform and the tails are not visible. TEM. Scale bars: A, B, 200 nm.

**Fig. 4.**
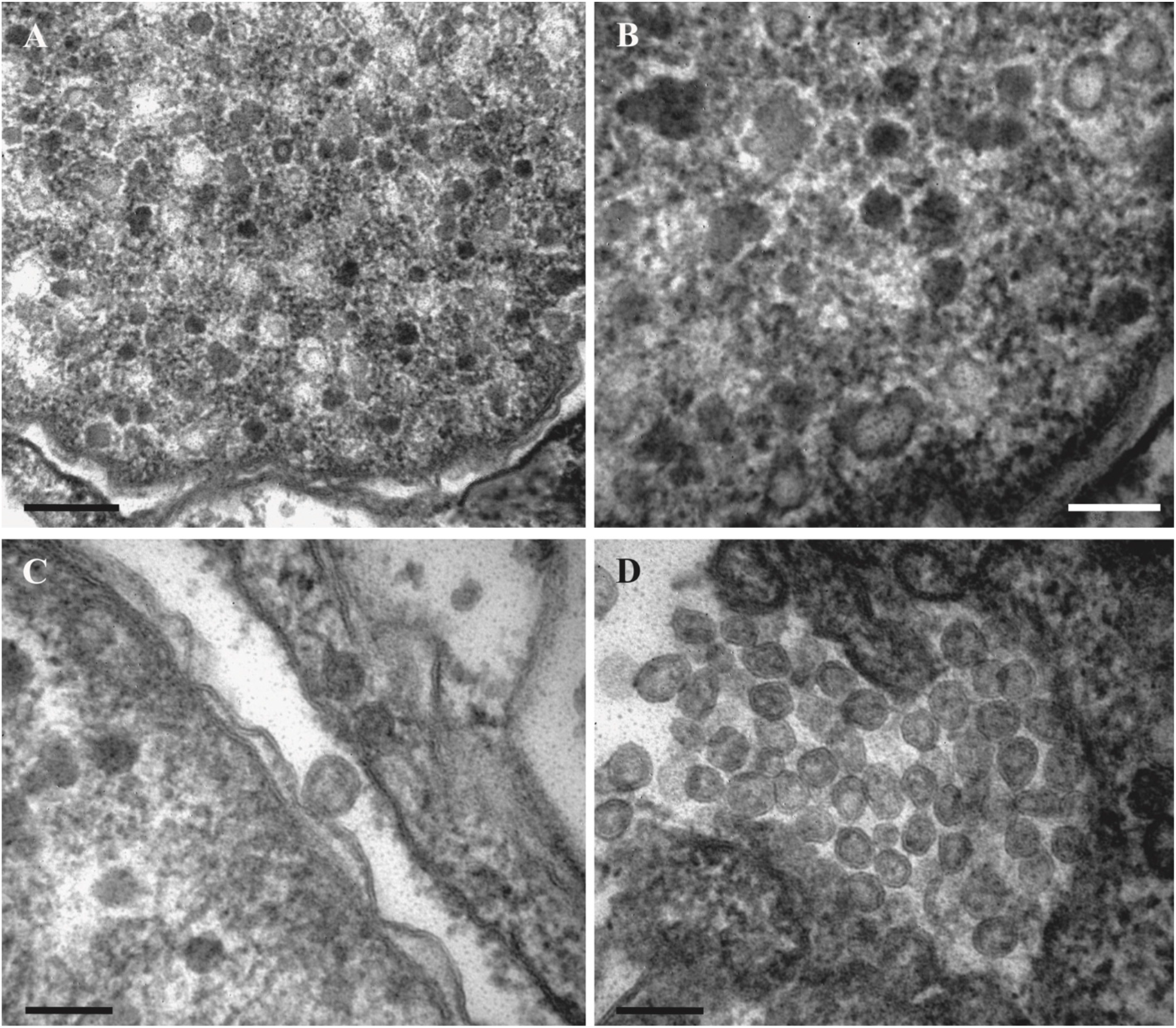
Virus-like particles associated with symbiotic bacteria in the tentacles of *Bugula neritina*. A, B. Electron-dense and electron-translucent VLPs inside the cytoplasm of the bacterial cell. C. VLP inside (left) and on the surface of the bacterial host. D, virions outside bacteria in and between epithelial cells of a bryozoan. TEM. Scale bars: A, 200 nm; B‒D, 100 nm.

### VLPs in bacterial symbionts of *Bugula neritina*

In *B. neritina* the virus-like particles were found in the symbiotic bacteria associated with the tentacle epidermis (Figs. 4, 6, 7) and those located inside the funicular bodies (Figs. 8, 9C, D, 10). In these two loci, bacteria were morphologically different, supposedly belonging to two different species. In both loci bacterial cells fall into three ‘ultrastructural types’, presumably representing the successive stages of bacterial transformation/destruction during the viral lytic cycle. The VLPs became visible in the bacteria of ‘types’ II and III.

#### VLPs and bacteria in the tentacles

The VLPs inside bacteria in the tentacle epidermal (ciliated) cells were predominantly oval and isometric, with diameters varying between 50 and 65 nm. Being either electron-dense or translucent, their content probably reflects different virion assembly intermediates (Fig. 4A, B). Some VLPs with relatively translucent content were detected on the surface of bacterial cells (Fig. 4C) or between them and bryozoan epidermal cells. They were either oval or polygonal with clearly recognizable thick ‘cover’/peripheral layer (Fig. 4D).

Bacteria of ‘type I’ represent non-altered cells with an ultrastructure typical for Gram-negative bacteria. These were coccoid or slightly elongated cells (Figs. 5, 6A) with a diameter/length of about 0.5-0.7 μm. A well-defined electron translucent nucleoid zone is surrounded by a thin peripheral layer of electron-dense cytoplasm enveloped by two membranes. Some of the cells were obviously dividing. Bacteria were recorded either directly in the cytoplasm or inside large vacuoles of the tentacle epidermal cells, individually or in groups. Some of them presumably occupied intercellular spaces. VLPs were not found inside or between the ‘type I’ bacterial cells.

**Fig. 5.**
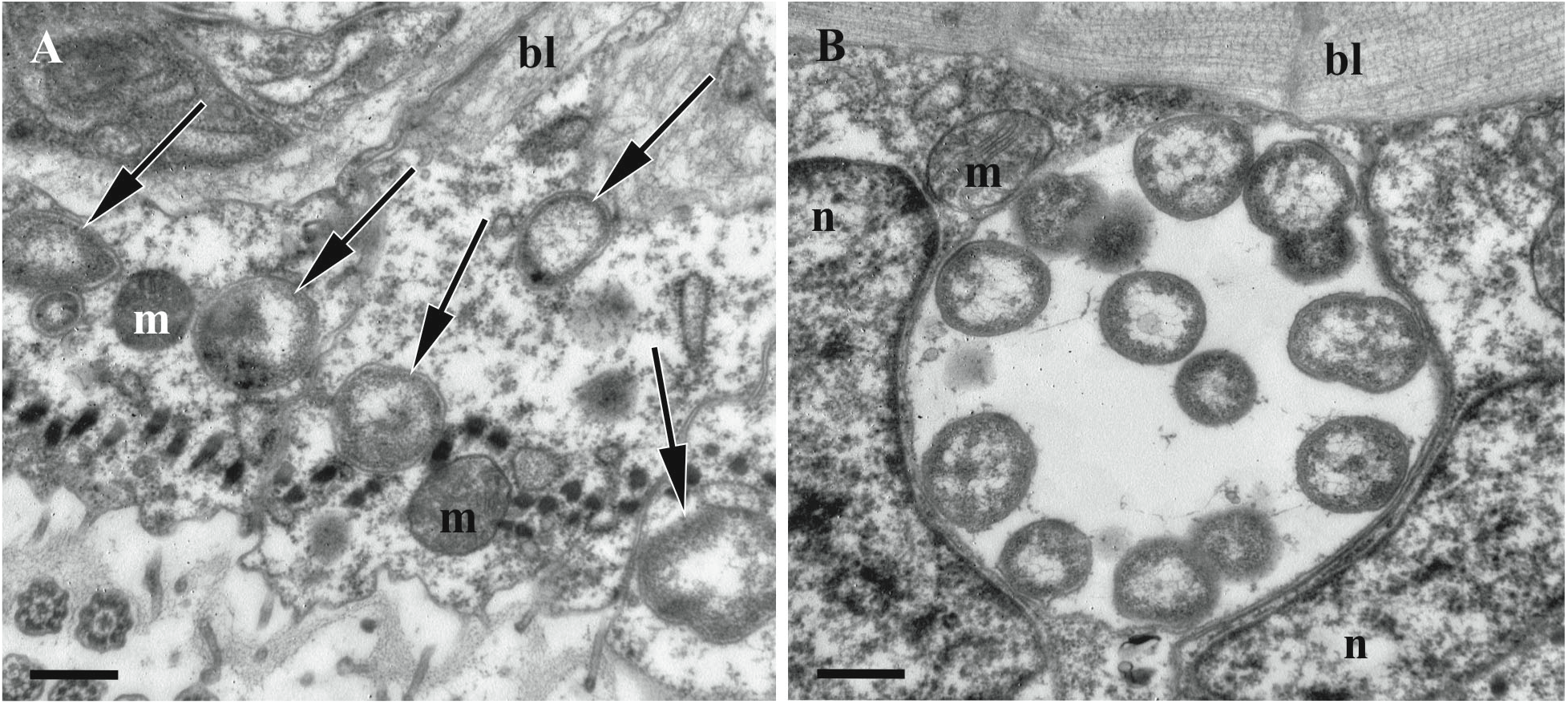
Coccoid symbiotic bacteria of ‘type I’ inside the cytoplasm (A, arrows) and a large vacuole (B) in the ciliated cells of the tentacles of *Bugula neritina* (sectioned kinetosomes and cilia are visible in the lower half of the image A). TEM. Abbreviations: bl, basal lamina; m, mitochondria; n, nucleus. Scale bars: A, B, 500 nm.

**Fig. 6.**
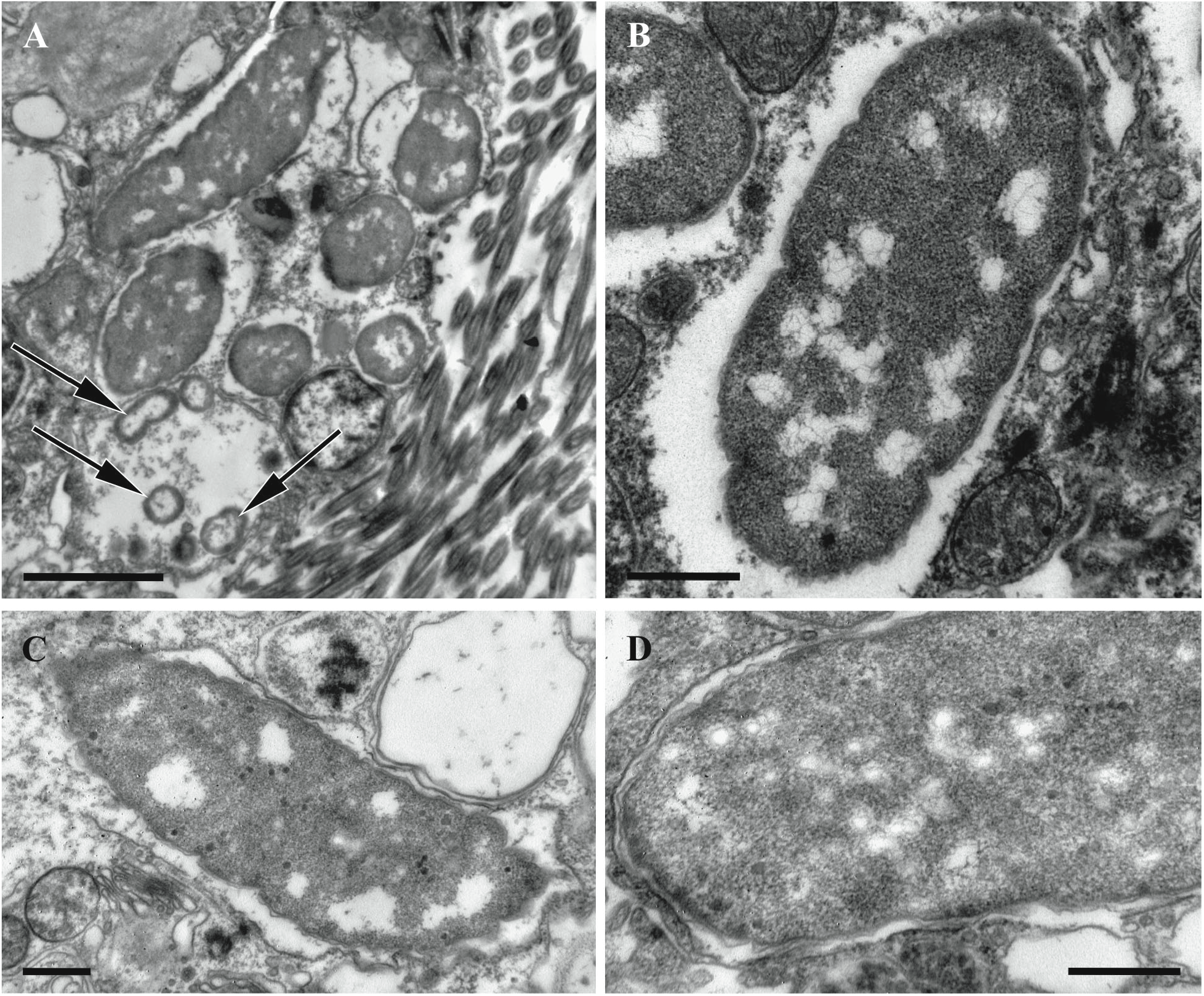
Symbiotic bacteria of ‘type I’ and ‘type II’ inside ciliated cells of *Bugula neritina* tentacles. A, groups of smaller coccoid bacteria of ‘type I’ (arrows) and the larger rod-like ‘type II’. B-D, ‘type II’ cells without (B) and with (C-D) few visible virus-like particles, both electron-dense and electron-translucent. TEM. Scale bars: A, 2 μm; B-D, 500 nm.

**Fig. 7.**
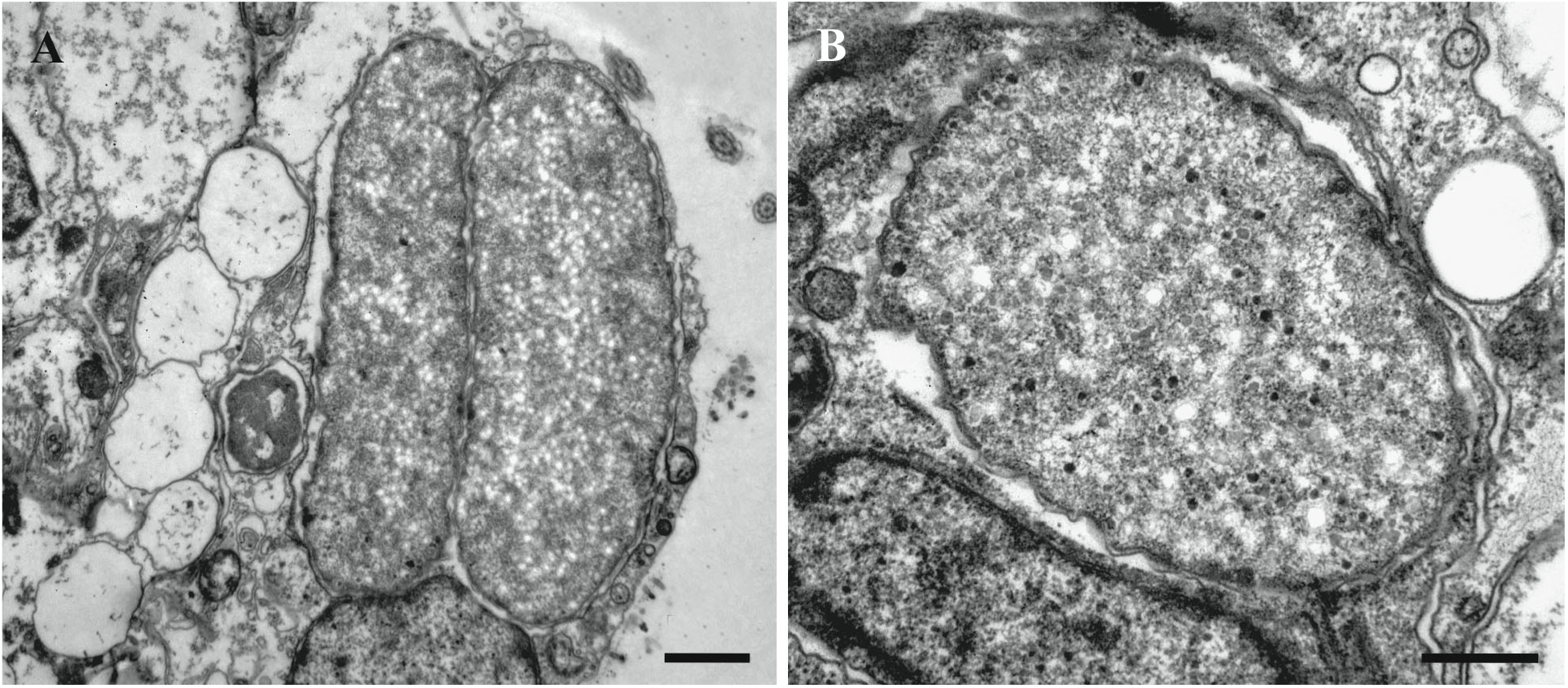
Symbiotic bacteria of ‘type III’ inside epidermal cells of *Bugula neritina* tentacles. A, B, bacterial cells with their cytoplasm full of viral particles, both electron-dense and electron-translucent. TEM. Scale bars: A, 1 μm; B, 500 nm.

**Fig. 8.**
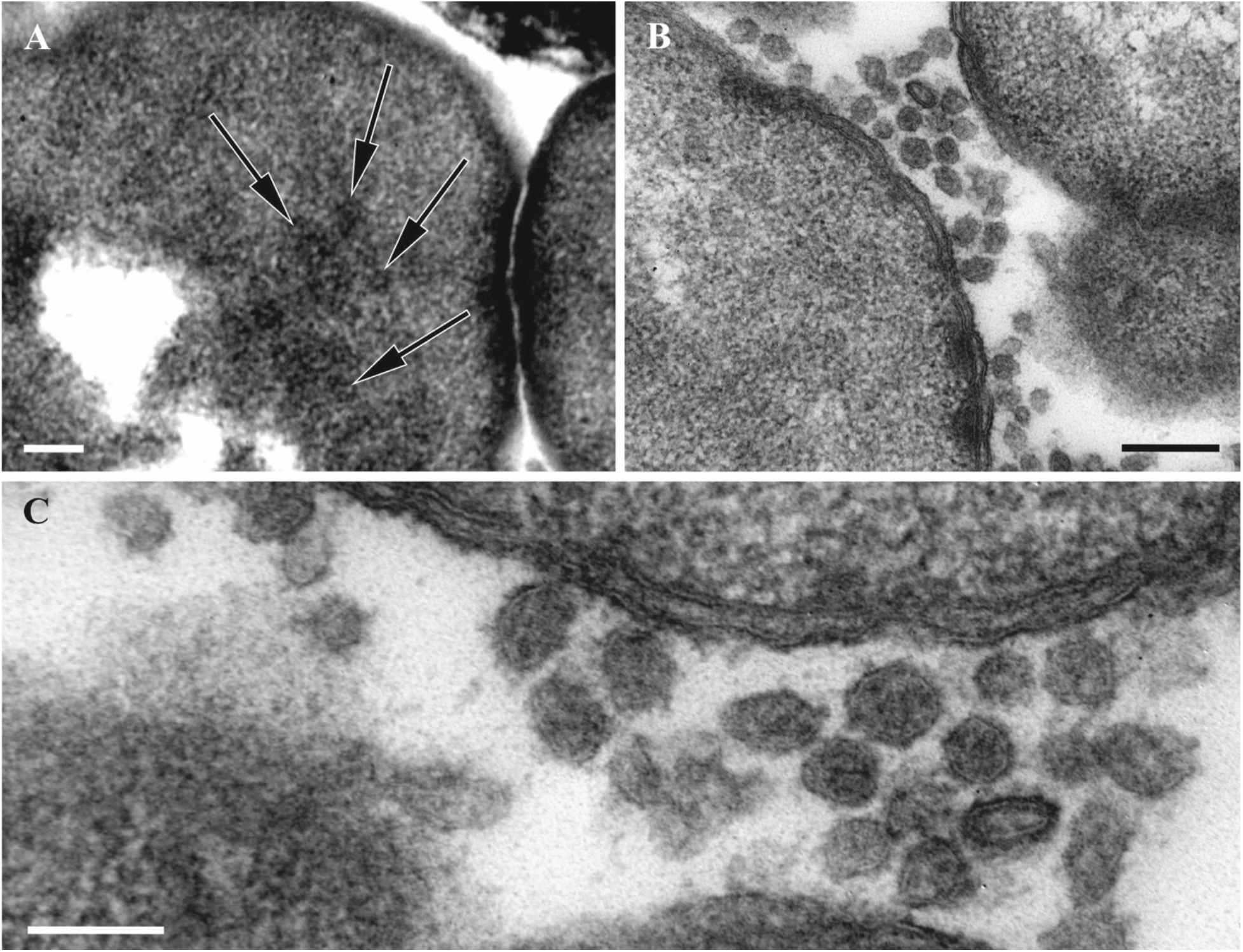
Virus-like particles inside (arrows) (A) and outside (B, C) of symbiotic bacteria in the funicular bodies of *Bugula neritina*. A, bacteria of ‘type II’. B, C, VLP between ‘type III’ bacteria. TEM. Scale bars: A, 500 nm; B, 200 nm; C, 100 nm.

**Fig. 9.**
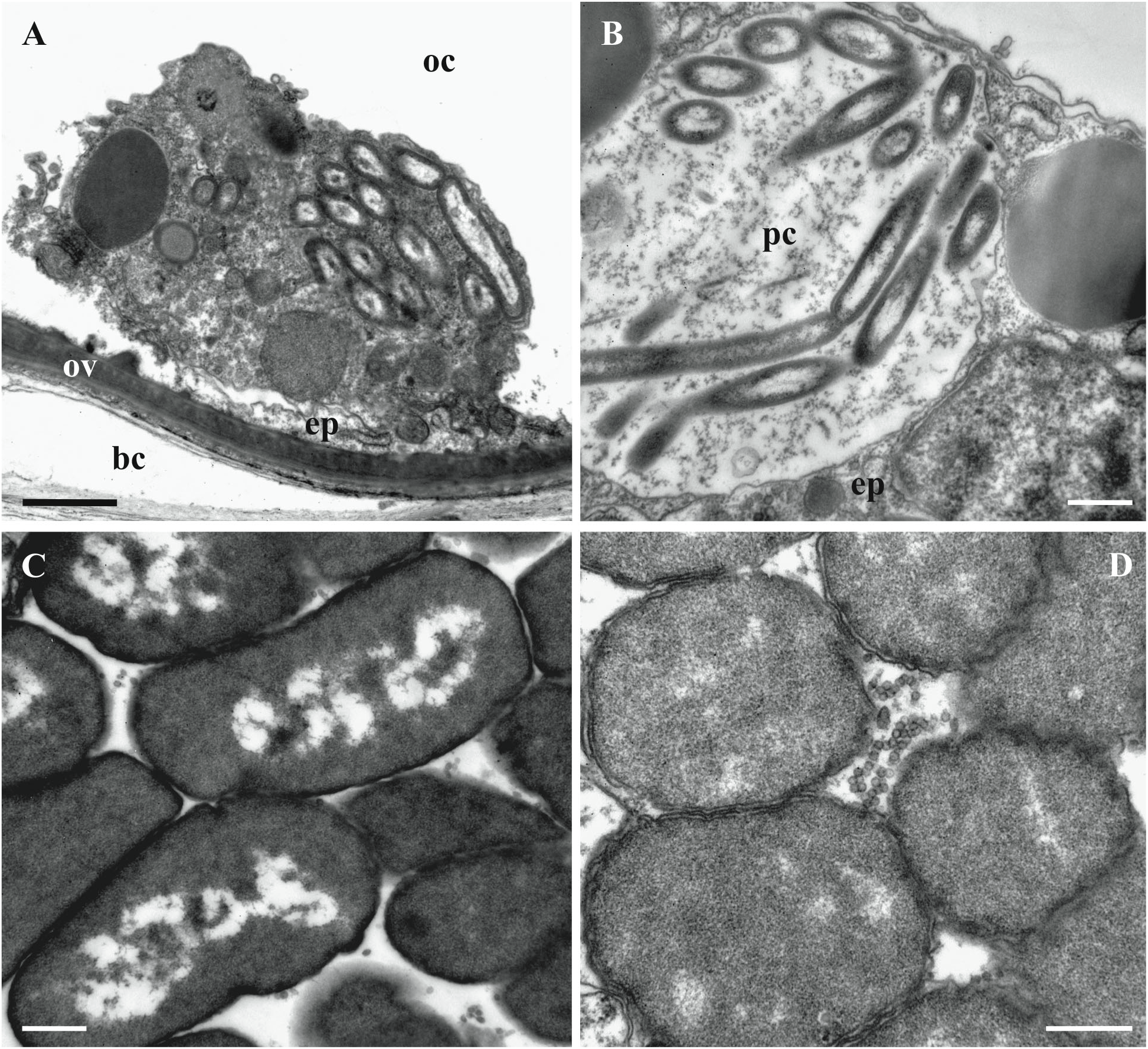
Symbiotic bacteria inside autozooids of *Bugula neritina*. A, B, intact bacteria of ‘type I’ in the cytoplasm of presumed amoebocyte (A) and peritoneal cell (B) of the body wall (ooecial vesicle). C, D, bacteria in the funicular bodies: ‘type II’ (C) and ‘type III’ (D) with VLP visible in between bacterial cells. TEM. Abbreviations: bc, incubation space of brood chamber; ep, epithelial cell; oc, coelom of ooecial vesicle; ov, wall of ooecial vesicle; pc, peritoneal cell. Scale bars: A, 1 μm; B-D, 500 nm.

**Fig. 10.**
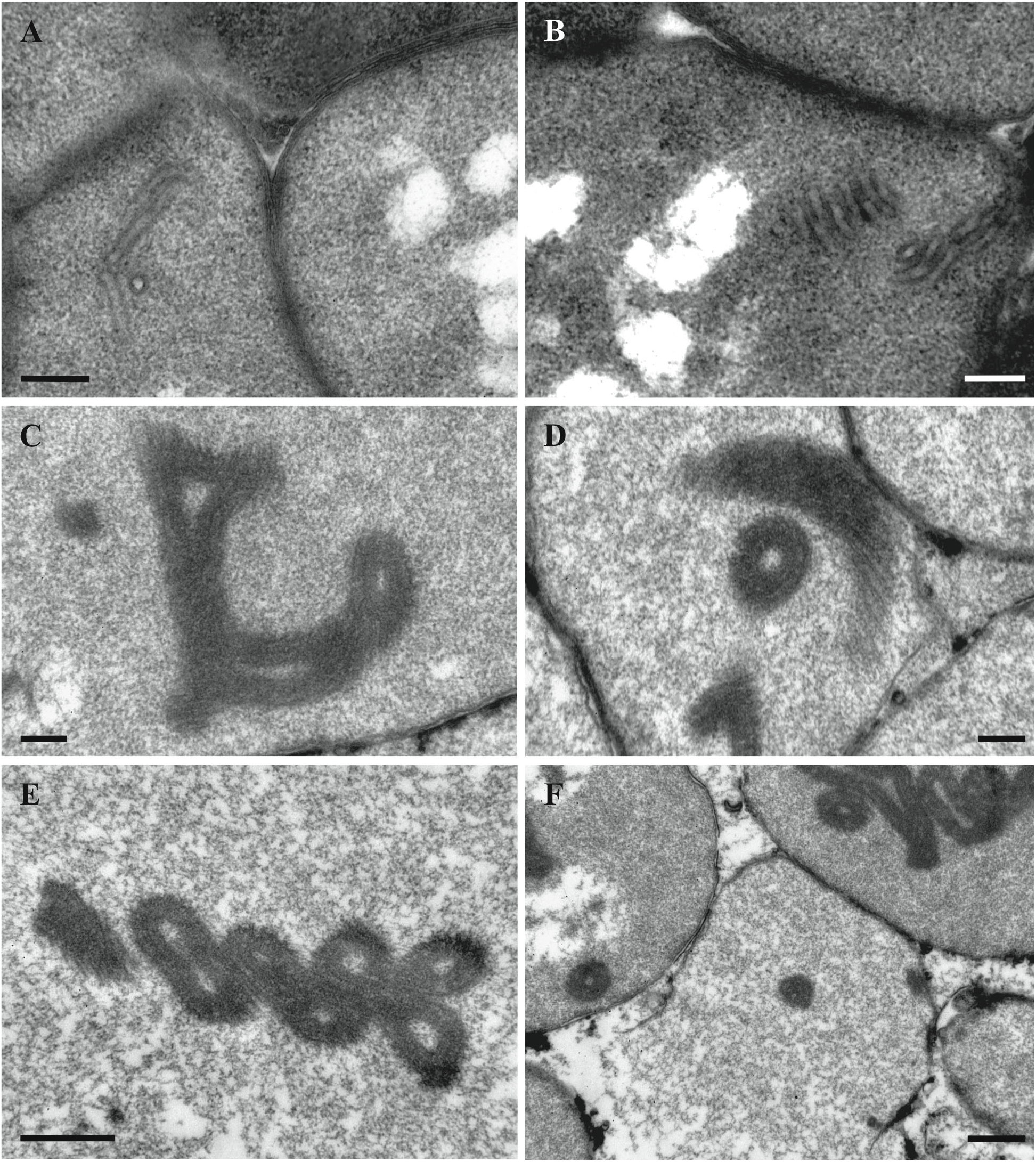
‘Tube’-like structures in the bacterial cells found in the funicular bodies of *Bugula neritina*. A, B, bacteria of ‘type II’ with tube’-like structures in the beginning of their formation. C-F, fully-formed ‘tube’-like structures in the bacteria of ‘type III’ and in destroyed bacteria (in F). TEM. Scale bars: A-D, 200 nm, E-F, 500 nm.

‘Type II’ bacterial cells (presumably next stage of their transformation) were rod-like and much larger than the coccoid form, reaching lengths of 4-5 μm and diameters of about 1.5 μm (Fig. 6). The cytoplasm was granular and electron-dense most of the volume, with the translucent nucleoid zone strongly reduced and fragmented. The cell wall was wavy (Fig. 6A, B). Some of these bacteria showed the presence of scant VLPs (Fig. 6C. D). The membranes of the vacuoles containing the ‘type II’ bacteria were often not clearly recognizable, possibly destroyed (Fig. 6A, B).

‘Type III’ bacteria were rod-like or ovoid. Their length was the same as in ‘type II’ cells, but the diameter increased up to 2-2.5 microns (Fig. 7). There were no traces of the nucleoid zone in ‘type III’ cells, and the cytoplasm was filled with abundant VLPs (Fig. 7).

The ‘type III’ bacterial cells were usually detected closer to the apical parts of the tentacles, and their numbers were significantly lower than the number of intact coccoid bacteria (‘type I’) distributed through the entire tentacle length.

#### VLPs and bacteria in the funicular bodies

The VLPs associated with the bacteria filling the funicular bodies (swollen parts of the funicular system, Fig. 2A, B) of *B. neritina* were oval-polygonal, isometric, with a capsid diameter of 50-70 nm. Most virions were detected in the spaces between bacterial cells (Figs. 8B, C, 9D), but individual VLP were also visible in the cytoplasm of bacteria (Fig. 8A).

As was the case in the tentacles, the three bacterial ‘ultrastructural types’ (but of presumably different species, see above) were also detected inside autozooids. Intact Gram-negative bacterial symbionts (‘type I’) were found in the epithelial cells (in the cytoplasm and inside vacuoles) of the introvert – the flexible part of the body wall everting and inverting during the tentacle expansions and retractions (see above), in the peritoneal cells of the ooecial vesicle wall and in presumed amoebocytes (Fig. 9A, B), i.e. the outfold of the body wall plugging the entrance to the ovicell (brood chamber). They were rod-like (length 2.0-2.5 μm and diameter 0.4 μm), had two outer membranes, a well-defined nucleoid zone and a thin layer of cytoplasm. They were present in groups in the cytoplasm as well as in the vacuoles. No VLPs were recognizable in them.

The ‘type II’ bacterial cells in the funicular bodies had a wide oval, often irregular shape (length up to 4.0-4.5 μm, width up to 1.5-1.7 μm) (Figs. 8A, 9C, 10A, B). Their cytoplasm was electron-dense and their nucleoid was fragmented (Fig. 9C). In some of these bacterial cells, ‘tube’-like structures 40-50 nm thick were detected in the cytoplasm, being separated or assembled into groups (Fig. 10A, B). The walls of these ‘tubes’ were morphologically and dimensionally similar to the membranes of the bacterial cell wall. In some cases these ‘tubes’ and cell membranes were connected.

The shape and size of the ‘type III’ bacterial cells (Figs. 9D, 10C-F) were about the same as in ‘type II’. The cytoplasm was relatively translucent and mostly homogeneously flocculent. Some of the ‘type III’ bacteria showed two well-recognizable, albeit deformed, cell membranes (Figs. 8B, C, 9D). Some of them, similar to the ‘type II’ bacteria (Fig. 10A, B), contained ‘tube’-like structures. In addition, only one cell membrane was often recognizable in the ‘type III’ bacteria that presumably were on the late stage of degradation (Fig. 10C-F). They contained large, twisted or curled electron-dense bodies with a spiral arrangement of parallel membranes. The bodies were up to several micrometers long, while their thicknesses varied, sometimes reaching 135 nm. The shape of these bodies strongly varied – we detected rings, U-shaped figures, complex multiple loops and other configurations. When the bacterial cells were destroyed, these bodies were released in the space between the bacteria.

Funicular bodies contained bacteria of either ‘type II’ or ‘type III’ separately, but the funicular bodies with different bacterial ‘types’ were detected in the same zooids.

### Virus-related structures in bacterial symbionts of *Paralicornia sinuosa*

In *P. sinuosa* the cytoplasm of morphologically altered bacterial cells found in the funicular cords (Fig. 2 C,D) contained spherical particles consisting of the cylindrical/tube-like elements evenly radiating from the central double-walled ‘core’ (Fig. 11). The diameter of these ‘sea urchin’-shaped structures was about 300 nm, while the individual cylinders/tubes were about 120 nm long and about 20 nm thick. In cross-section the core with the bi-layered darker periphery (core wall?) and lighter central parts were recognizable at high magnification (Fig. 11B and insert). The cylinders/tubes were apparently interconnected by the electron-dense layer slightly below their distal tips. The proximal ends of the tubes also appear connected, forming a spherical ‘double-walled’ core of the particle that has a diameter of 50-60 nm. In the medial sections of the particles about 20-21 cylinders/tubes are visible, leading to an estimated 400-450 of them in one particle. The particle diameter seems to be stable/identical, suggesting that the size and/or the number of the cylinders/tube-like elements is tightly controlled during particle development.

**Fig. 11.**
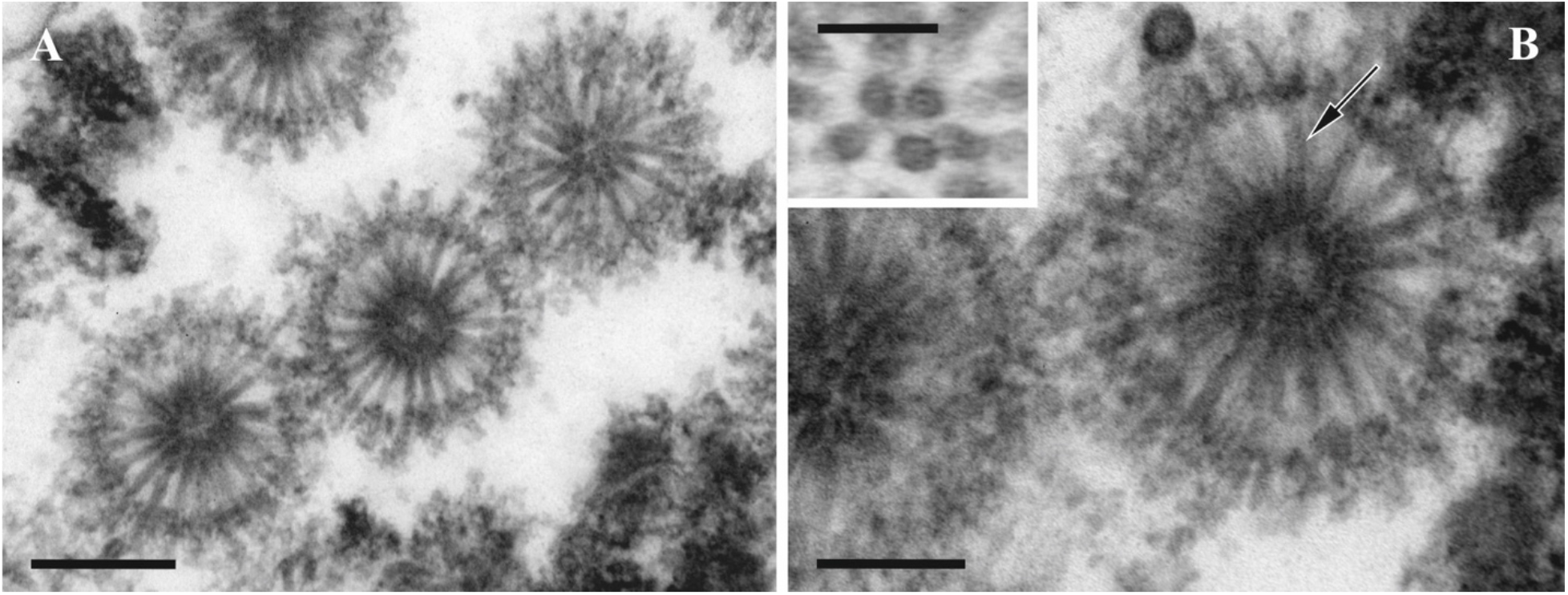
A, B. Structures presumably derived from viruses in bacterial symbionts of *Paralicornia sinuosa*. Insert: cross-section of radial cylinders showing darker periphery and lighter central part (also shown by arrow in B) with a dark central ‘spot’. TEM. Scale bars: A, 200 nm; B, 100 nm; insert, 50 nm.

Similar to *Bugula neritina*, closer examination of the morphology of bacterial cells also revealed three different ‘ultrastructural types’, which we again interpret as corresponding to the stages of the virus-like particle development before bacterial lysis. ‘Type I’ – presumably intact or slightly altered Gram-negative bacteria. Such cells were rod-like, 3-4 μm long and about 1 μm wide (Fig. 12A, insert) The central part of the cell is occupied by filaments of nucleoid surrounded by an electron-translucent zone that often included electron-dense areas of various size and shape. The peripheral cytoplasm is electron-dense, granular, without inclusions, surrounded by a plasma membrane that was poorly recognizable in some cases.

**Fig. 12.**
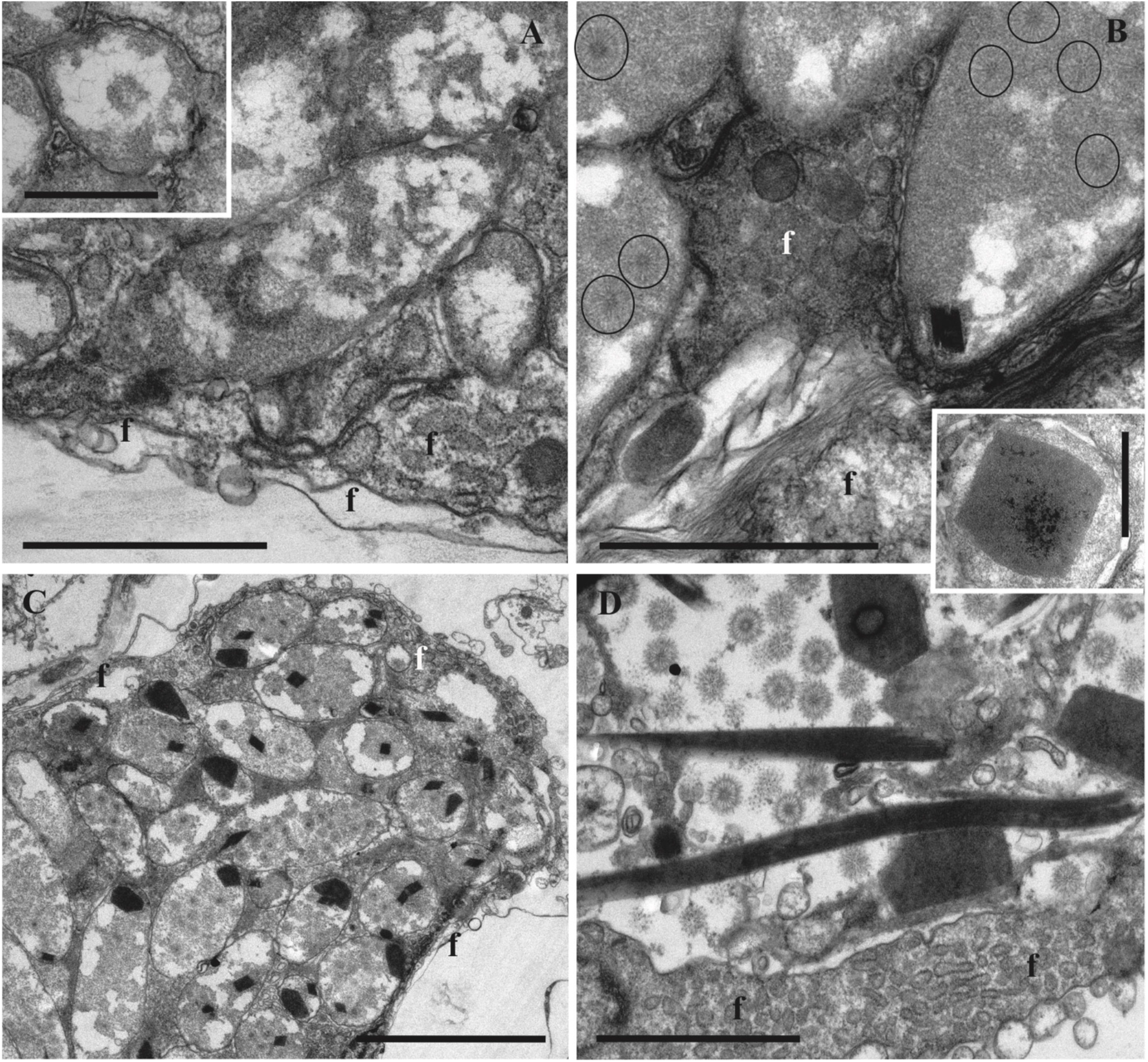
Various stages in the lytic cycle of bacteriophages in the symbiotic bacteria of *Paralicornia sinuosa*. A and insert, longitudinal and cross-sections of non-/less altered bacteria (‘type I’ cells) inside funicular cord. B, VLP (in circles) becoming visible inside bacteria predominantly filled with electron-dense cytoplasm (‘type II’ cells), nucleoid is not recognizable; insert shows paracrystalline body at higher magnification. C, bacteria (‘type III’ cells) filled with viruses and paracrystalline structures. D, bacteriophages and paracrystals inside the cavity of the funicular cord after bacterial destruction. TEM. Abbreviations: f, cells of funicular cord. Scale bars: A, B, D, 2 μm, inserts, 1 μm; C, 5 μm.

Bacterial cells of ‘type II’ presented moderate alterations in the cell ultrastructure presumably corresponding to the initial phase of VLP development. Cell length and width were increased, sometimes doubled (up to 8 and 2 μm, respectively), and the cell wall often took on a wavy appearance. The nucleoid fibrils disappeared, and most of the cytoplasm became homogeneously electron-dense. The viral particles and paracrystalline structures became visible inside the bacteria (Fig. 12B). In ‘type III’ cells the VLPs filled most of the cell volume (Fig. 12C). The paracrystalline bodies became more numerous, acquiring very different shapes, from irregular polygons to long rods with a length of up to 8-10 μm (Figs. 2C, D, 12C, D). At high magnification the densely packed parallel fibrils were visible in them (Fig. 12B, insert).

After destruction of bacteria, both the viral particles and paracrystals were freely distributed inside the cavity of the funicular cords. In some occasions we observed them in the zooidal coelom too, obviously following disintegration of the funicular cord wall.

## Discussion

Detection of new viruses is often a serendipitous event during ultrastructural studies (e.g. Reuter 1975; Vacelet & Gallissian 1978; Crespo-González et al. 2008). In our case we found VLPs when studied of symbiotic associations of bacteria with an invertebrate. We explain the different bacterial ‘ultrastructural types’ detected in the tissues of two bryozoan species as stages in the gradual destruction of bacterial cells caused by the virus development.

The two variants of VLPs found in the symbiotic bacteria of *Bugula neritina* (compare Figs. 4 and 8) were similar to the capsids of tailed bacteriophages. No tails were observed, however. We suggest that the tail appendages are probably present but indiscernible in the TEM-images, as, for example, in the case of podoviruses with their short tails. Nonetheless, we cannot exclude that the VLPs belong to non-tailed phages that possess the virions of similar size such as, e.g. in the family Tectiviridae (Mäntynen et al. 2019).

Our observations indicate that the phages of symbiotic bacteria of *B. neritina* act as pathogenic agents, destroying bacteria in the course of their multiplication in the lytic cycle.

The situation observed in bacteria from the funicular system of *B. neritina* is special. We suggest that the ‘tubes’ visible in the cells of the second ‘morphotype’ are invaginations of membranes of the bacterial cell wall, and can later cluster as the bacteria are lysed, forming complex structures in the cytoplasm. Subsequently, elongation and spiral twisting of these ‘tubes’ could result in the formation of flexible ‘rods’. Further curling of these rods yields structures of various shapes. A similar effect is observed in the action of lactocins and of the viruses on Gram-positive bacteria of the genus *Lactobacillus* (Cuozzo et al. 2003; Chibani-Chennoufi et al. 2004).

Bacteria discovered in the tentacles and the funicular system of *B. neritina* have a different morphology (coccoid *vs* rod-like), suggesting different (and independent) ways of their acquisition. The former potentially could penetrate into tentacle tissues directly from the surrounding water. This kind of interaction between bryozoan hosts, bacteria and phages might be common, but more extensive ultrastructural research on bryozoan tentacles is required to show whether this mechanism of infection is widespread. A supporting argument in this case is the discovery of similar symbionts and phage particles in the tentacles of *B. neritina* collected in the Mediterranean Sea on the coast of Spain (Vishnyakov, Schwaha, Souto, Ostrovsky, unpublished data).

The symbiotic bacteria occupying funicular bodies in the bryozoan zooidal cavity have been identified as Candidatus Endobugula sertula and are vertically transmitted through the larva and are thus inherited by the first zooid (ancestrula) during metamorphosis (Haygood & Davidson 1997; Sharp et al. 2007a). Zooidal budding then results in colony formation and the spread of the symbionts through it. Occupying the transport funicular system and utilizing the nutrients from it, the symbionts multiply, triggering the formation of the funicular bodies (Karagodina et al. 2018). Bacteria partly stay in these bodies and partly move to the ooecial vesicle of the brood chamber by an unknown mechanism. Woollacott and Zimmer (1975) showed the presence of bacteria inside the lacunae of the funicular cords of this species, which could be the pathway for the symbionts traveling through zooids. Our data indicate an opportunity for bacterial transport to the ooecial vesicle by amoebocytes, but more research is needed to evaluate this hypothesis.

The available data are insufficient to conclude whether the VLP production in symbiotic bacteria results from infection by externally acquired bacteriophage particles or is due to induction of the prophages or prophage-like elements present in the genomes of these bacteria. The infection hypothesis seems more plausible for bacteria found in *B. neritina* tentacles, but the scenario potentially could be different in case of the funicular body symbionts. We suggest that in this species the phages circulate in the bryozoan host population together with bacteria by vertical transfer, which does not require repeated external infection. Those symbionts that remain in the funicular bodies finally collapse due to viral activity. In contrast, some bacteria move by an unknown mechanism from the funicular bodies towards the brood chamber containing phages in the form of prophages in the bacterial genome. We speculate that the superficially intact ‘type I’ bacteria we found in the peritoneal cells and presumed amoebocytes in the ooecial vesicle are, in fact, infected with prophages. Similar aggregations of the “tiny dark granules” (possibly bacteria) were earlier described and/or illustrated inside or close to the ooecial vesicle in *B. neritina* (Mathew et al. 2018, p. 8) and a related bryozoan species (Ostrovsky et al. 2009; Ostrovsky 2013a, b) using histological sections. The reconstructed draft genome of Ca. *E. sertula*, consisting of 112 contigs, from a metagenomic assembly did not uncover any prophage-like elements (Miller et al 2016a); however, it is possible that such elements were absent in those particular specimens (one pooled collection of B. neritina larvae and several autozooids and ovicells of one *B. neritina* colony), or that these prophages were not recovered by the assembly algorithms. Interestingly, many short contigs (~2 kbp) were recovered from the metagenome (Miller et al 2016b), and some of these contigs might represent viral sequences.

The next step is a transfer of infected bacteria from the ooecial vesicle through its wall to the brood cavity. Moosbrugger with co-authors (2012) showed the presence of bacteria inside an ovicell containing an embryo (see also Sharp et al. 2007a).

Accordingly, if our model is correct, prophages are transferred to the brood chamber with their bacterial hosts finally reaching the larvae. This enables them to be transmitted to the next bryozoan generation. Potentially, the phages and bacteria found in the tentacles might circulate in the host in a similar way, but it then remains unclear how they could reach larvae in the brood chamber. The only imaginable strategy is their transfer during oviposition when the tentacle crown delivers the zygote to the brood cavity (its mechanism reviewed in Ostrovsky & Porter 2011 and Ostrovsky 2013a). This way, however, appears less probable.

The induction hypothesis appears to be the only possibility to explain the formation of the structures observed in *Paralicornia sinuosa.* These structures do not resemble any known bacteriophages and appear to be too large to be interpreted as the virions of a novel bacteriophage family. At the same time the formation of these structures is clearly associated with cell destruction by lysis. We speculate that the particles observed in *P. sinuosa* symbionts are similar to the metamorphosis associated complexes (MAC) recently discovered in the free-living bacteria *Pseudomonas luteoviolaceae* (Shikuma et al. 2014). MACs are encoded by prophage-like elements in the *P. luteoviolacea* genome, and their formation is associated with host cell death and lysis. Structurally, MAC are an assemblage of multiple contractile systems related to the contractile tails of myoviruses (tailed bacteriophages with contractile tails) (reviewed by Leiman et al. 2009 and Taylor et al 2018). The tail-like structures in MAC are assembled in a sea urchin-like pattern that fills almost the entire host cell. The baseplates of the individual contractile systems point outwards and are interconnected by a network formed by the homologs of the tail fibers (Shikuma et al. 2014). Interestingly, the formation of MAC in *P. luteoviolacea was* associated with the formation of 2D paracrystalline arrays that were tentatively interpreted as the crystals of the sheath protein (Shikuma et al. 2014). In *P. sinuosa* bacterial cells, the sea-urchin like structures are also associated with large protein crystals (Fig. 12), adding to the similarity of these two systems. Although the formation of similar crystalline bodies was previously reported in certain cells infected by viruses (Lawrence et al., 2014), the physiology of this process and the possible function of these structures are poorly understood.

The MACs of *P. luteoviolacea* function to deliver a protein signal into the cells of the larvae of the tubeworm *Hydroides elegans.* The delivery takes place when the larva contacts the biofilm containing the MACs. This causes the contraction of the tail-like structures and the penetration of the tail tube tips into the animal cells (Ericson et al. 2019). The delivery of the protein signal induces larval metamorphosis (Shikuma et al. 2014; Ericson et al. 2019). It is currently unclear whether bacteria can benefit from this effect.

The features of the structures that we detected in *P. sinuosa* symbionts are compatible with a presumed MAC-like nature. The length of their cylindrical elements is about 120 nm, i.e. comparable to the phage tail of the bacteriophage T4 contractile tail (114 nm long) (Ackermann 2009). Although no data are currently available on a potential involvement of MAC-related structures in interspecies communications between bacterial symbionts and their *P. sinuosa* host, it seems a plausible hypothesis. In this regard it is important that small (compared to *P. luteoviolaceae* MAC) particles released by the lysed symbiont cells inside bryozoan host may move from its funicular system to the zooidal coelomic cavity and further to the surrounding water via coelomopores (Ostrovsky & Porter 2011).

Interestingly, the onset of the MAC or VLP production appears to occur almost simultaneously in multiple bacterial cells within one funicular body (Figs. 9C, D, 12). This suggests the triggering of induction by some cue generated by bacteria or by the animal host.

We speculate that maintaining an excess of symbiotic bacteria in the funicular bodies becomes energetically disadvantageous at a certain point, and viruses destroying bacteria thus act as regulators of their numbers. Such phages are termed mutualistic, and their participation in indirectly regulating the number of symbiotic bacteria and processes carried out by symbionts has been described for aphids and some other insects (Moran et al. 2005; Bordenstein et al. 2006; Weldon, Oliver, 2016). This relationship as also been assumed for the symbiotic associations of cnidarians with their microbial symbionts (Rohwer Thurber 2009; Roossinck 2011).

Our study is the first step towards future research on viruses of the bacterial symbionts of Bryozoa. The next step should include determination of the systematic status of both the viruses and their bacterial hosts. The wide range of bioactive substances identified from bryozoans (reviewed in Sharp et al. 2007b, see also Maltseva et al. 2016) suggests a wide distribution of symbionts inside these invertebrates. These symbionts, in turn, could host various viruses – but this research has just begun.

## ACKNOWLEDGEMENTS

This study was performed using the laboratories and equipment of the Centre for Molecular and Cell Technologies, Saint Petersburg State University. Dr K. Tilbrook, Oxford University, kindly helped during collecting and identification of the Australian material. Drs A. Hoggett and L. Vail, Lizard Island Research Station, Australian Museum, kindly provided all the necessary help during field work on the Great Barrier Reef. We thank Dr M. Stachowitsch, University of Vienna, for linguistically revising the early draft of the manuscript. We are also deeply indebted to anonymous reviewers who helped to improve the manuscript. Planning of the research, TEM microscopy, data processing and analysis, and manuscript preparation were performed at the Saint Petersburg State University, being funded by the Russian Science Foundation (grant 18-14-00086). The animal samples for microscopy were processed at the facilities of the University of Vienna. The *E. coli* samples processing and a part of the data analysis were performed at the Winogradsky Institute of Microbiology, Research Centre “Biotechnology” of the Russian Academy of Sciences.

## COMPLIANCE WITH ETHICAL STANDARDS

### ETHICAL APPROVAL

No ethical issues were raised during our research.

### SAMPLING AND FIELD STUDIES

No permits for sampling and field studies were required.

